# Effects of Alzheimer’s disease and formalin fixation on the different mineralised-iron forms in the human brain

**DOI:** 10.1101/2020.06.02.129593

**Authors:** Louise van der Weerd, Anton Lefering, Andrew Webb, Ramon Egli, Lucia Bossoni

## Abstract

Iron accumulation in the brain is a phenomenon common to many neurodegenerative diseases, perhaps most notably Alzheimer’s disease (AD).

We present here magnetic analyses of post-mortem brain tissue of patients who had severe Alzheimer’s disease, and compare the results with those from healthy controls. Isothermal remanent magnetization experiments were performed to assess the extent to which different magnetic carriers are affected by AD pathology and formalin fixation.

While Alzheimer’s brain material did not show higher levels of magnetite/maghemite nanoparticles than corresponding controls, the ferrihydrite mineral, known to be found within the core of ferritin proteins and hemosiderin aggregates, almost doubled in concentration in patients with Alzheimer’s pathology, strengthening the conclusions of our previous studies. As part of this study, we also investigated the effects of sample preparation, by performing experiments on frozen tissue as well as tissue which had been fixed in formalin for a period of five months. Our results showed that the two different preparations did not critically affect the concentration of magnetic carriers in brain tissue, as observable by SQUID magnetometry.

## Introduction

Mineralized iron in the form of magnetite was first detected in the human brain almost thirty years ago^1^. Since then, magnetite has been detected in multiple organs, including the heart^2^, the liver^2^ and, most recently, the cervical skin^3^. These findings are of biological interest because magnetite, an iron-oxide mineral hosting both Fe(III) and Fe(II) in its crystal structure, has been proposed as a source of cellular toxicity, involving many different mechanisms. For example, in addition to the well known hypothesis that iron may produce reactive oxygen species via the catalytic Fenton reaction^4, 5^, a more recent hypothesis concerns the effects of the magnetic field associated with the magnetic moment of these nanoparticles: a few studies have related the prolonged effect of such local static magnetic fields to a series of harmful biological phenomena, including protein structural alteration^6^ and suppression of the threshold of the neuronal action potential^7^.

Recent literature has suggested a relation between the presence of magnetite nanoparticles in the brain and the incidence of Alzheimer’s pathology and its hallmark, amyloid beta plaques^8–11^. In particular, one study has reported higher levels of magnetite in the temporal cortex of Alzheimer’s patients when compared to controls^12^. Therefore, being able to determine the magnetic properties of iron nanoparticles, their concentration in the brain, their source, i.e. whether they are of biogenic or of anthropogenic origin^13^, and their reactivity in physiological conditions^14, 15^ may reveal important information on Alzheimer’s disease and possibly other neurodegenerative diseases that are accompanied by brain iron accumulation^4^.

While the most compelling evidence for the presence of magnetite in the human material has come from High Resolution Transmission Electron Microscopy (HRTEM)^1^ and X-ray techniques^16^, magnetic analyses of Isothermal Remanent Magnetization (IRM) are often employed as complementary tools, or as a means of reporting concentrations of magnetite nanoparticles in the tissue^2, 3, 13, 17–19^. Although these latter measurements require very sensitive magnetometers, they offer the ability to study tissue blocks larger than those that can be studied by TEM, and the dimensions of such blocks are comparable to the spatial resolution of in vivo imaging techniques such as magnetic resonance imaging (MRI), meaning that magnetometry can potentially be employed to guide the interpretation of the contrast of, for example, magnetic susceptibility-weighted MRI images. Moreover, magnetometry requires minimal sample preparation, contrary to TEM, thus lowering the risk of sample contamination.

The observation that IRM curves obtained from human material saturate at magnetic fields of 300 mT or lower, and at temperatures between 50 K and 293 K^2, 3, 13, 17, 19, 20^, and the occasional observation of the Verwey transition^3, 19^, support the use of magnetometry techniques to detect and potentially quantify magnetite and/or oxidised magnetite, i.e. maghemite, nanoparticles in post-mortem or ex-vivo human samples.

Furthermore, extending the IRM measurements below 10 K offers the possibility to detect residual magnetization from ultra-fine magnetite/maghemite particles, if present, and from ferrihydrite nanoparticles that are found in the core of ferritin^19^ and hemosiderin proteins^4, 21^.

Despite the increasing use of magnetometry for the study of brain material, we note that sample preparation is very different in published studies, and it is important to determine whether measurements are dependent or independent of such methods. Some authors have used formalin-fixed tissue^17, 18, 20^, while others have used fresh-frozen tissue^1, 3, 13, 19^. The main argument put forward against formalin-fixation is the potential leaching of iron from the tissue. However, there are only very limited studies on the effects that fixation and storage may have on the magnetic properties of the tissue^22, 23^. Some researchers therefore propose to only use frozen materials. However, formalin-fixed brain material is much more widely available, and offers the possibility to perform multi-technique studies on the same specimen^20^.

In this study we focus on two types of magnetic carriers detectable by magnetometry, in the temporal cortex of severe Alzheimer’s patients, i.e. magnetite/maghemite and ferrihydrite. In particular, we aim to determine which iron-oxide minerals are distributed in the brain tissue, and whether they all present the same coercivity field, using IRM curves. Additionally, we aim to validate and extend previous results in which we reported higher ferrihydrite levels in the temporal cortex of AD patients, and unaltered levels of magnetite/maghemite in presence of AD. Finally, to determine whether formalin fixed samples are valid for this purpose, we assessed whether formalin fixation (for a period of approximately five months) had an effect on the concentration of the magnetic carriers in the tissue. Fresh-frozen material from healthy controls was used as the gold standard, and the effects of relatively short formalin fixation and Alzheimer’s disease were investigated against this standard.

## Materials and Methods

### Tissue selection

Frozen (−80 °C) post-mortem brain tissue blocks from the medial temporal gyrus of patients with severe AD (N=9) and age- and gender-matched controls (N=9) were obtained from the Netherlands Brain Bank (NBB) and the Normal Aging Brain Collection Amsterdam (NABCA). The autopsy, fixation and storage protocols of these two collections were identical. All donors gave written informed consent for the use of their tissue and medical records for research purposes. The AD cases were selected based on pathological reports (Braak stage 5 or higher) and the appearance of hypo-intense bands and/or foci on the T_2_*-weighted MRI images of the contralateral hemisphere of the same subject^24^. Details of the subjects’ demographic are found in Table 1.

**Table 1.**
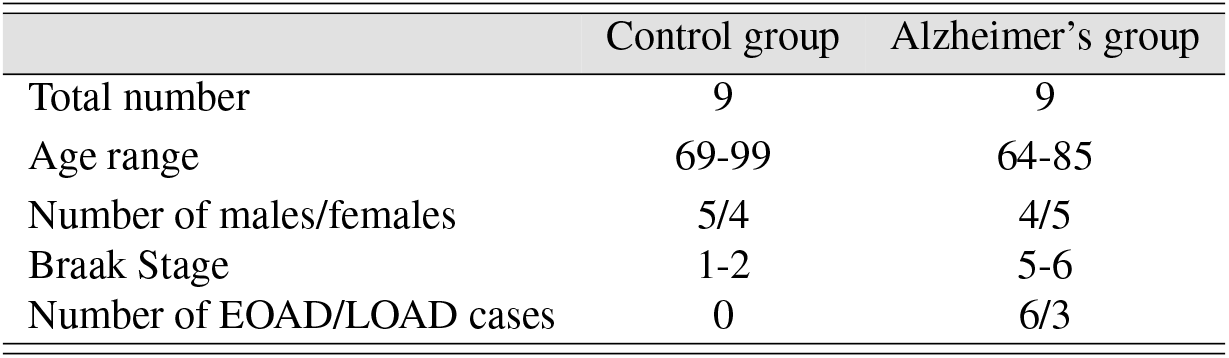
Demographic of the subjects in this study. EOAD refers to Early Onset Alzheimer’s disease. LOAD refers to Late Onset Alzheimer’s disease.

Each tissue block contained predominantly grey matter. The blocks were cut, while frozen, into smaller cubes of approximately 2 mm^3^ each; the sections were randomized and then pooled back into two paired groups: the first was immediately freeze-dried and prepared for SQUID magnetometry, while the second one was immersed in fresh formalin (4 % formaldehyde, DiaPath), and kept at 4 °C for a period of approximately five months before being removed from formalin and freeze-dried. The pH of the storing formalin, after five months of storage (and up to one year), did not drop below 7.0. Care was taken not to contaminate the tissue with metal objects. The tissue was sealed in sterile plastic tubes before being prepared for the magnetic analyses.

### SQUID magnetometry experiments and data analysis

The dried tissue was pelleted with the use of two non-metallic pistons, placed in the center of the SQUID-straw and kept in position by two symmetrically bent straws from each side, as in Ref.^25^. This method was employed to prevent the use of gel capsules that occasionally show remanent magnetization. The sample was then loaded in a Quantum Design MPMS-XL SQUID magnetometer with the reciprocating sample option (RSO), and centered in the apparatus while at room temperature. The field was then quenched (via a magnet reset). This is the most reliable and time-saving manner to minimize offset fields due to flux creep in the magnet coil^26, 27^. The offset field (−2 Oe) was characterized on a Pd reference sample. However, residual fields in the magnet coil depend on the magnetic history of the superconductor, and vary as a function of the initially-applied field. Since quenching the magnet before measuring each data-point was not possible, in this study we did not exceed fields of 1 T, to avoid introducing a bias in the saturation field.

Two IRM curves were acquired: one at 100 K and a second one at 5 K. Both temperatures were reached in zero-field cooled condition. The acquisition protocol was as follows: a magnetic field was applied and allowed to stabilize for 150 s before being set to zero, and the residual magnetic moment was measured after a waiting time of 180 s. The measurement occurred over a range of 3 cm of sample movement, data were averaged over 3 scans with 5 oscillations each at 1 Hz, and an iterative regression routine was used to fit the raw data.

IRM raw curves were empirically fitted to the Langevin model, as described previously^20^: the fit was weighted by the experimental uncertainty on each data-point with a custom-written MATLAB script. An offset parameter was added to account for non-reproducible residual fields that would yield a non-demagnetized initial state. The saturation remanent magnetization (SIRM) was extracted from the fit. In contrast to our previous work^18^, here we do not analyze the mean magnetic moment, since its interpretation can be confounded by particle’s interaction, as it will be discussed later.

Finally, we note that the SIRM values refer to the dry weight of the tissue. The wet-to-dry weight ratio (R_*wd*_) for the AD samples was 5.9±0.2, which was very similar to that of the healthy controls: R_*wd*_ = 6.2±0.2. These numbers are in agreement with a previous study which reported no difference in R_*wd*_ between AD and controls^28^, and when formalin-fixed tissue is used^29^, since formalin fixation does not alter the total water content of the tissue^30^. However, it is worth noting that the white matter contains consistently less, i.e. almost 50 %, water than the gray matter^29^.

### Statistical analysis

To test whether SIRM values differed between control and AD subjects, we performed a Mann-Whitney U-test (two-sided). The small number of samples and the non-normal distribution of the SIRM values justify the use of non-parametric tests. The p-level used to refute the null hypothesis was 0.05.

To test whether formalin fixation affected the IRM values at saturation, the Wilcoxon signed rank test (two-sided) was performed between the data obtained from the frozen and the formalin-fixed batch (paired test). In addition, a Bland-Altman plot was constructed. The statistical analysis was done in Origin 2018b.

Finally, a cluster analysis was performed on the 100 K-IRM data obtained from the frozen samples batch. First, the offsets of raw IRM curves that started from an incomplete demagnetized state were corrected. Curves containing more than one outlier data-point were excluded from the analysis. Second, the remaining curves were normalized by the mean remanent magnetic moment acquired between 70 and 120 mT. Saturation-normalized curves (NIRM) obtained in this manner reflect the intrinsic magnetic properties of the particles, regardless of their concentration. NIRM curves were pooled into two clusters using the Mathematica *FindClusters* function with a *k*-mean clustering algorithm. The two clusters differ by their median acquisition fields (H_*m*_), defined as the field required to acquire half of the saturation remanence identified with the mean moment in the 70-120 mT range. Cluster 1 (C_1_) appears rather heterogeneous, however, increasing the total number of clusters causes a further partition of C_1_ into additional clusters which differ only with respect to their noise characteristics.

Due to the large measurement noise, H_*m*_ values were estimated by fitting individual NIRM curves with a phenomenological relation:

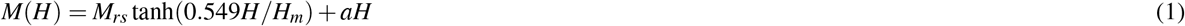

where M_*rs*_ is the saturation remanence of the low-coercivity component, H_*m*_ the corresponding median acquisition field, and *aH* a linear term accounting for the contribution of a heavy saturation tail or a distinct, high-coercivity component (Fig. 1). Cluster analysis was not performed on the IRM data obtained at 5 K because these curves are heavily under-saturated at 1 T.

**Figure 1.**
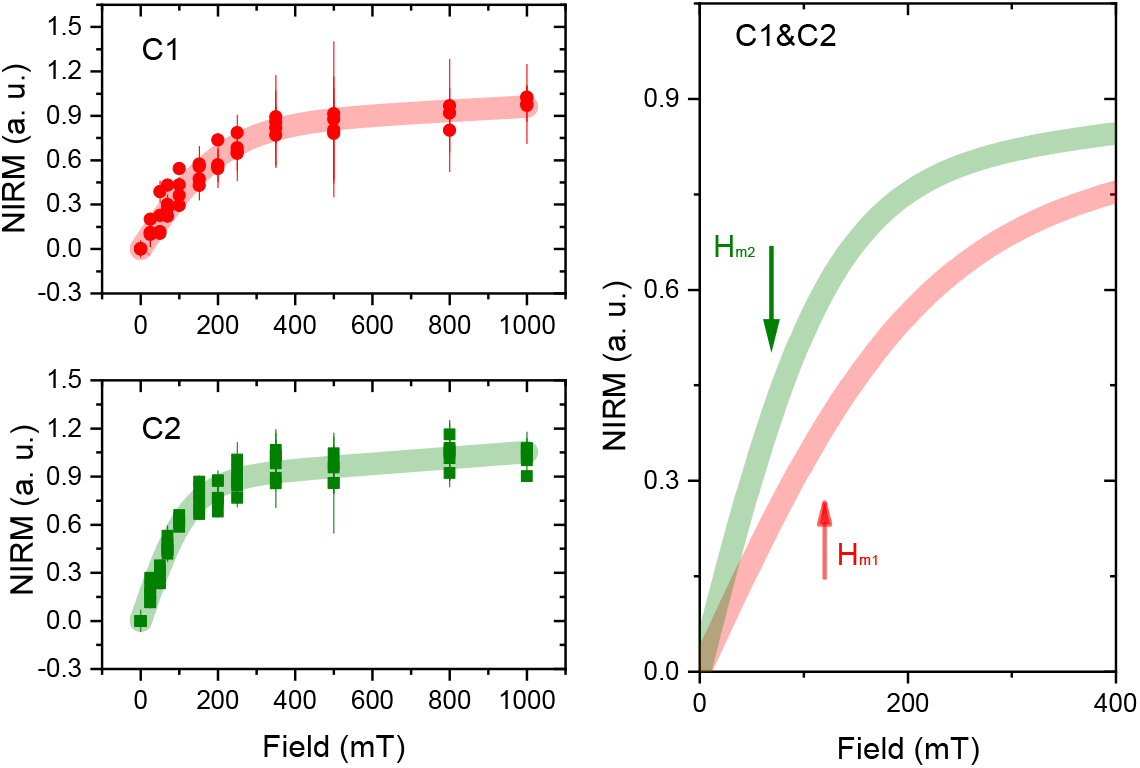
Result of the cluster analysis on the normalized IRM curves (NIRM). NIRM data on the samples included in the analysis for cluster 1 and 2 (top and bottom left panels) are shown with red/green data points. The experimental error is also displayed. The red/green trace overlaid on the data-points represent the global fit to the experimental data of the corresponding cluster. In the right panel, only the fitted curves are displayed, and are zoomed, to highlight the difference in the median acquisition fields, which are marked by solid arrows. The best fit parameters were H_*m*_=120.3 mT, a= 2.3 × 10^−4^, M_*rs*_= 0.79, for C_1_ and H_*m*_=70.8 mT, a= 1.7 × 10^−4^, M_*rs*_= 0.84, for C_2_.

## Data availability

Data generated in this study and the SQUID experimental sequence are available upon request from the corresponding author in .txt format.

## Results

In this study, we performed IRM measurements on human brain tissue, which can be modelled as an ensemble of magnetic nanoparticles embedded in a non-magnetic matrix. IRM measurements on an ensemble of nanoparticles can probe the irreversible switch of the magnetic moments of the particles induced by the application of an external magnetic field, once the blocked state has been established. At the measurement temperature of 100 K, maghemite nanoparticles of approximate size of 14 nm, or larger, will contribute to the IRM signal^31^. This is an estimate based on the assumption that magnetite nanoparticles are completely oxidised in the form of maghemite, and that the characteristic time *τ*_0_ is equal to 10^−9^ s and the anisotropy constant *K*=2.6 × 10^5^ erg/cm^3^. Given that earlier HRTEM studies have identified magnetite nanoparticles in brain tissue with a median size of 20 nm^13^, the vast majority of the maghemite/magnetite nanoparticles in the tissue should therefore contribute to the IRM signal at 100 K.

The magnetic properties of the brain samples measured in this study are summarized in Table 2. These data show that the magnetic moment measured at 100 K is one order of magnitude above the noise floor of the SQUID magnetometer. However, the relatively low SNR at this temperature (in the range of 1-10), together with the large standard deviation demonstrate how challenging these measurements are. Therefore, extracting biologically-relevant information from these type of measurements should be done carefully.

**Table 2.** Summary of the magnetic properties of the brain samples measured in this study. ‘AD’ refers to Alzheimer’s disease group and ‘HC’ to the healthy control group.

| T = 100 K<br>($10^{-10}$ Am<sup>2</sup>) | frozen samples | | fixed samples | |
| --- | --- | --- | --- | --- |
|  | AD | HC | AD | HC |
| mean saturated magnetic moment | 5.35 | 3.43 | 4.87 | 4.69 |
| median saturated magnetic moment | 4.65 | 3.10 | 4.58 | 3.53 |
| standard deviation saturated magnetic moment | 3.95 | 1.28 | 2.44 | 4.05 |

| T = 5 K<br>($10^{-9}$ Am<sup>2</sup>) | frozen samples | | fixed samples | |
| --- | --- | --- | --- | --- |
|  | AD | HC | AD | HC |
| mean saturated magnetic moment | 36.95 | 20.11 | 35.38 | 11.50 |
| median saturated magnetic moment | 44.14 | 19.77 | 29.14 | 12.19 |
| standard deviation saturated magnetic moment | 14.74 | 12.48 | 18.66 | 3.42 |
| mean dry mass (mg) | 63.6 | 58.2 | 51.4 | 41.4 |

All the brain samples in this study showed a low-coercivity component which is completely, or partially, saturated at 300 mT. This suggests the presence of ferrimagnetic maghemite or magnetite. To gain more understanding of the specific iron-oxide mineral under study, we performed a cluster analysis on the IRM curves, as discussed above.

The normalized SIRM curves at 100 K (SIRM_100*K*_) obtained from the frozen samples belong to two clusters, characterized by different median acquisition fields H_*m*1_= 120±27 mT (C_1_) and H_*m*2_=70±15 mT (C_2_) (for NIRM data and fit see Fig. 1). This parameter can be used as a proxy for the *speciation* of the iron-oxide mineral, as will be discussed in the next section. Interestingly, we found that C_1_ contains predominantly control subjects (one AD vs four HC), while C_2_ contains predominantly AD cases (four AD vs two HC). We remind that some NIRM curve/subjects were excluded from the analysis, as discussed above.

The IRM curves obtained at 5 K did not show saturation up to 1 T, and indeed from our previous studies, saturation was not observed up to 6 T^20^. This is an indication of the presence of high coercivity components, that we ascribe to ferrihydrite: the mineral predominantly found in the core of ferritin proteins^21^. However, we do not exclude that the 5K-IRM signal also originates from other iron minerals such as hematite, and goethite^21^, with high coercivitites, and/or smaller magnetite nanoparticles (of about 5 nm minimum size), with smaller coercivities.

In addition to the analysis of the saturation field, it is illustrative to analyse the value at which the IRM curves saturate (SIRM), as the latter is related to the concentration of the magnetic carrier responsible for that saturation, as it will be discussed in more detail in the next section. SIRM data in the AD male group reported higher values than the AD female group: the mean SIRM at 100 K for the male AD group was more than two times higher than the one for the female group, while at 5 K the male AD group showed 1.3 times higher SIRM than the female paired group (Fig. 2). We note that the age distribution between AD males and females subjects was the same. These gender differences were not statistically significant, probably due to the small sample size. We note that this gender difference was not found in the healthy control group.

**Figure 2.**
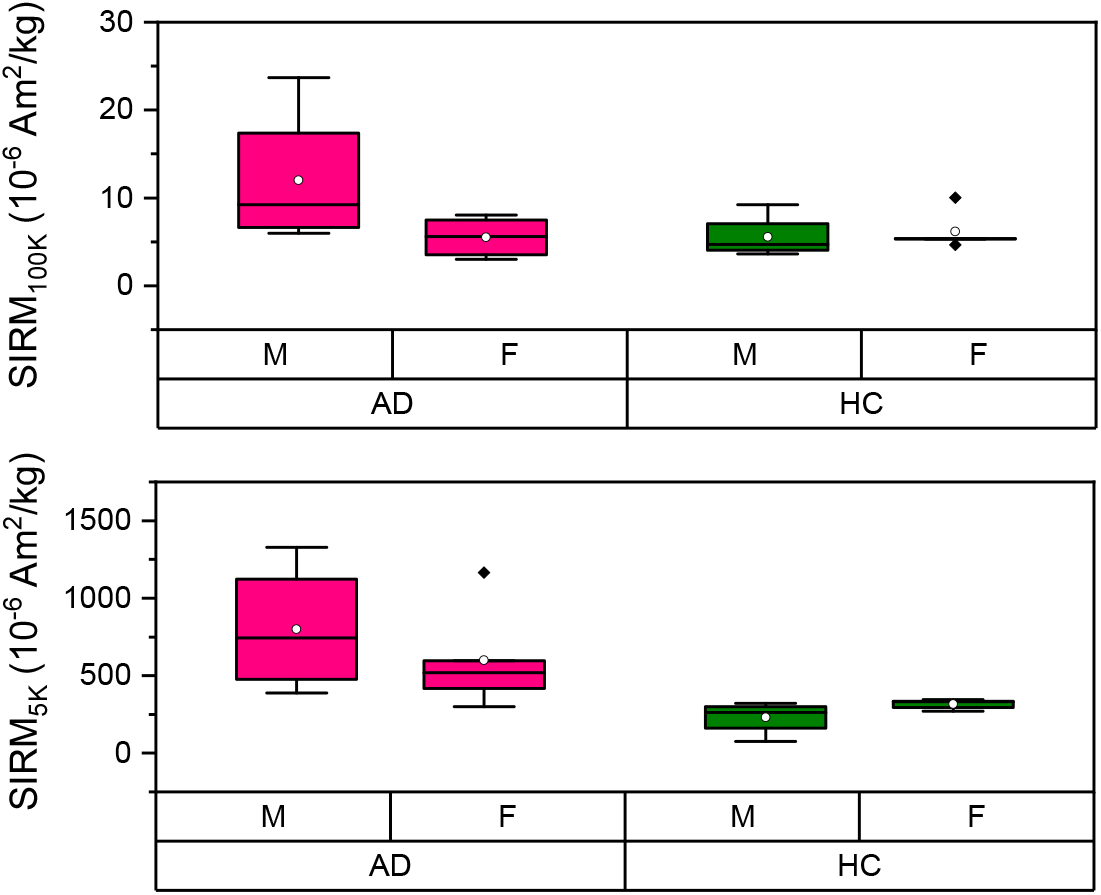
Box-plot of the SIRM data at 100 K (top panel) and 5 K (bottom panel) stratified by gender and diagnosis. Data were obtained from the frozen-samples batch. The pink box-plots refer to the diseased cases (AD), while the green box-plots refer to the healthy controls (HC). ‘M’ and ‘F’ refer to the ‘male’ and ‘female’ group, respectively.

When the SIRM data were stratified by diagnosis, no difference in the SIRM_100*K*_ was found between the control and AD group, while the SIRM_5*K*_ values were significantly higher in the AD than the control group (p=0.0015), as reported in Fig. 3.

**Figure 3.**
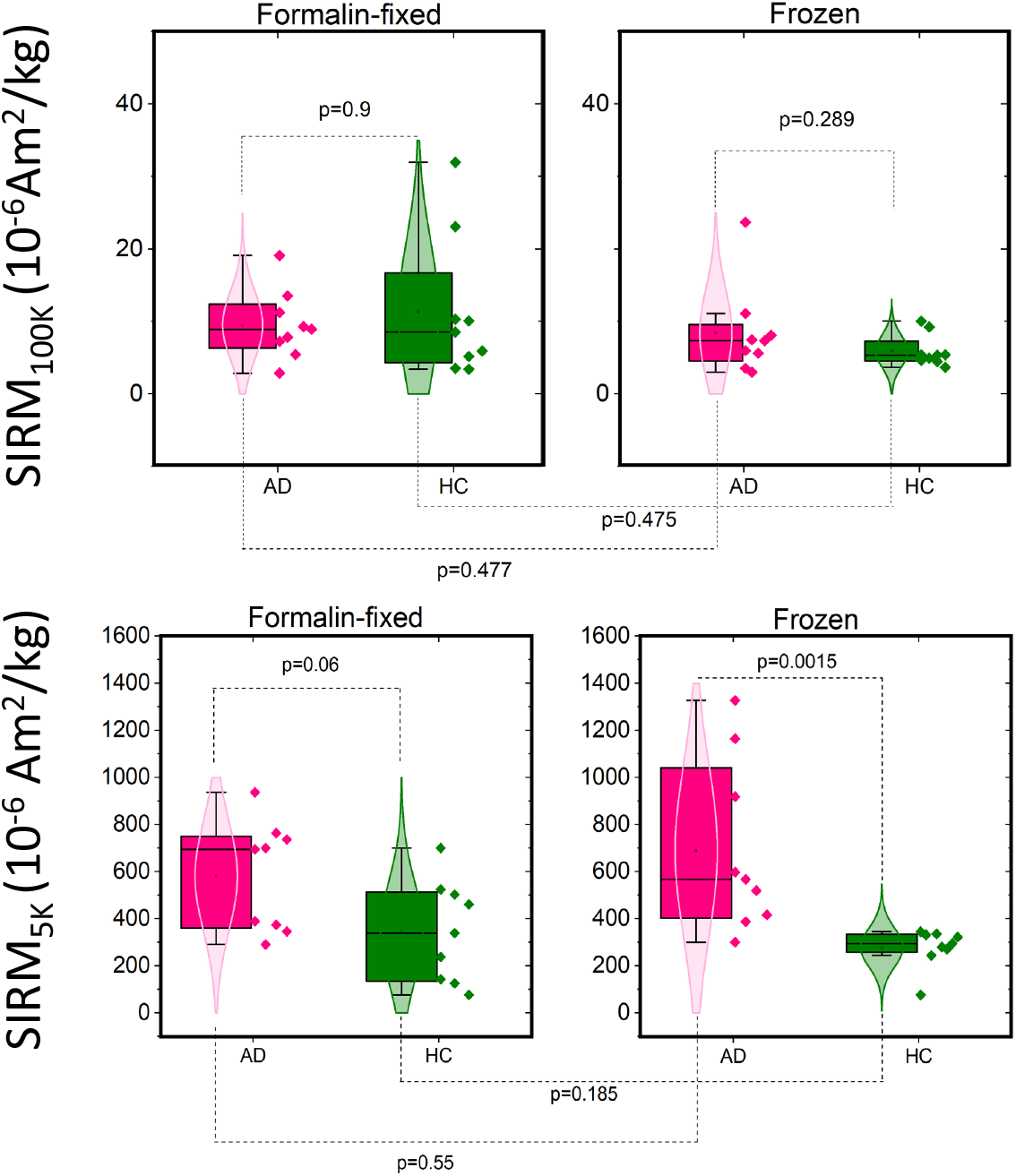
Box and scatter plots of the SIRM values summarizing the effect of formalin fixation and disease. (Top) SIRM measured at 100 K, on the frozen (right) and formalin-fixed (left) batches. The data points are individual samples. The p-value on top of the data refers to the Mann-Whitney U-test, while the p-value between the same group-type but different sample preparation (bottom lines) refers to the Wilcoxon signed paired test (bottom). (Bottom) SIRM measured at 5 K, on the frozen (right) and formalin-fixed (left) batches. Green color is used for the healthy control group and pink is used for the AD group.

To test for the effect of formalin fixation on magnetic remanence, magnetometry experiments were performed on paired formalin-fixed samples, as explained previously. The results of the Wilcoxon-signed paired test are reported in Fig. 3, and show no systematic deviation between the two groups of samples at either 100 K or 5 K. However we note that the p-value of the Mann Whitney U-test for the SIRM_5*K*_ values dropped to p=0.06 for the formalin-fixed batch.

In order to exclude systematic differences in SIRM values between frozen and formalin-fixed material, we constructed a Bland-Altman plot (Fig. 4) taking the frozen-sample data as the reference value. A positive difference in SIRM values would correspond to an apparent loss in magnetic carriers, while a negative one would indicate an apparent gain. The vast majority of the samples fell within the limits of agreement and no proportionality bias was found for either 100 K or 5K data. The 100 K data show a relatively large negative bias (−3.25) when compared with mean SIRM values, while the 5 K data show a positive bias (19.83). In both cases, the bias was not significant, since the line of equality is within the confidence interval. Finally, upon visual inspection of the plot, it can be noticed that the variability of the data does not appear homogeneous across the plot: data showing the smallest mean SIRM values, particularly at 100 K, also display small variability, while the scatter increases with increasing mean SIRM.

**Figure 4.**
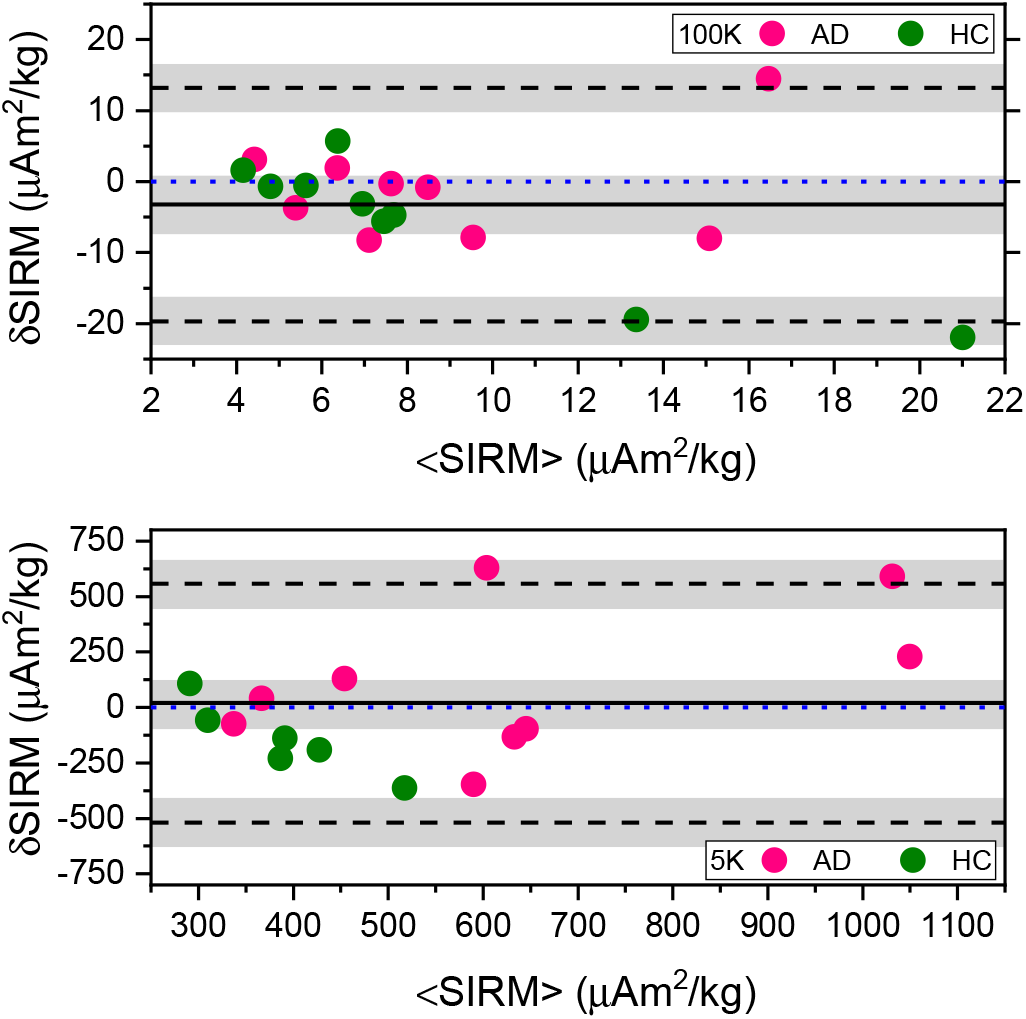
Bland-Altman plot of the difference in SIRM values before and after fixation. Differences in SIRM values (*δ*SIRM) obtained at 100 K (top row) and at 5 K (bottom row) are plotted against mean SIRM values (<SIRM>). The solid line represents the bias, while the dashed black lines represent the limits of agreement. The blue-dotted line represents the line of equality. The gray rectangles present confidence interval limits for mean and agreement limits. The green color is used for the healthy controls (HC) and pink is used for the Alzheimer’s cases (AD). Linear regression was used to exclude for proportionality bias. At the 5% level, the data slope was not different from zero.

## Discussion

This study was performed on paired frozen and formalin-fixed samples obtained from the post-mortem medial temporal gyrus of Alzheimer’s patients and healthy control subjects, in which IRM experiments were performed at two temperatures. We found, in all the subjects, measurable concentrations of magnetite nanoparticles and ferrihydrite nanoparticles. No difference in the concentration of the former mineral was found between the groups, while the concentration of the latter was significantly elevated in the AD subjects. Formalin fixation of the tissue taken from the same subject and same region did not have a significant effect on the magnetometry results. However, as we argue below, this may be due to a lack of sensitivity of the technique.

The analysis of the coercive field of remanence of the IRM curves provides an indication of the iron-oxide or oxyhydroxide mineral which is contributing to the signal. The median acquisition fields of 70 mT and 120 mT correspond to approximate coercive fields of H_*c*1_ ~ 47 mT and H_*c*2_ ~ 80 mT, if the median acquisition field is identified with the coercivity of remanence (H_*cr*_), and if a ratio H_*c*_/H_*cr*_ =1.5, for single-domain particles^32^ is used. These coercive fields are slightly larger than expected for partially-oxydised spherical single-domain magnetite nanoparticles^33^. These higher-than-expected coercivity values could be explained by the presence of elongated particles, which would result in a higher shape anisotropy. However, this hypothesis is not supported by TEM analysis of magnetic material extracted from brain tissue, and in-situ imaging^3, 13^. An alternative explanation could be the presence of single-domain particles that are magnetically interacting, as suggested by previous experiments on brain^22^ and dermal material^3^, which reported a Wohlfarth’s ratio < 0.5. Magnetostatic interactions increase the field required to magnetize a particle cluster (e.g. by step-wise IRM acquisition, as in our case) with respect to the field required to demagnetize it. In this case, H_*c*_ values depend heavily on the inter-particle distance, and the overall cluster geometry^32^. Therefore, with our current data, we can state that the observed coercive fields are generally in agreement with magnetite grains and partially oxidised maghemite nanoparticles, but we cannot unequivocally confirm that the magnetic grains in our samples are single-domain and interact with one another.

Moreover, the fact that the median acquisition fields cluster into two groups suggests that two different populations of magnetite particles may be present in the brain, as suggested by TEM and EELS experiments^13^, and that possibly one type could dominate in the AD group.

The absolute value at which the IRM curve saturates is proportional to the number of maghemite/magnetite particles per sample unit mass, the proportionality constant being the saturation magnetization M_*rs*_, which ranges from ~ 40 Am^2^/kg to 76 Am^2^/kg, for maghemite particles of size varying from 14 nm to bulk^34^. The actual M_*rs*_ value depends on the particles size^35, 36^, shape, and surface anisotropy, which cannot be obtained from the current data. Therefore, we performed the statistical analysis directly on the SIRM values.

Our analysis suggests the presence of a gender difference in SIRM_100*K*_ and SIRM_5*K*_ values, which is present only in the AD group. This result differs from that observed in an earlier magnetometry study^12^. Although there is not enough statistical power here to draw strong conclusions about a gender effect, we note that in a recent study, cerebellum iron-levels were elevated in males compared to females, only in a group with ‘high (AD) pathology’. However, the authors failed to found the same difference in the inferior temporal cortex which, on the other hand, reported iron levels correlating with antecedent cognitive decline in those individuals who had underlying plaque and tangle pathology^37^.

*Total* iron levels are known to increase in different brain areas of AD cases^28^, with the temporal cortex being one of the most and earliest affected regions^24, 38^. It is not yet clear what the underlying phenomena leading to brain iron accumulation in the context of AD are. However, it is likely that localized accumulation of iron might deprive other brain areas of this metal, thus impairing neuronal function^5^. Altered iron-homeostasis is reflected by the modified heavy-to-light (H/L) ratio of the chains composing the shell of ferritin proteins: AD brains present increased H/L ratio in the frontal cortex, as a possible means of copying with increased iron levels^39^. The increase in SIRM_5*K*_ values in the AD group found in our work is in agreement with previous observations of increased total and ferritin-iron levels in the AD subjects^40^, as assessed by different techniques such as histology^41, 42^ and MRI^43, 44^, and confirms that iron homeostasis is heavily altered in specific brain regions of AD patients.

Although iron levels are clearly increased in AD, we did not find any difference in the SIRM values/concentration of magnetite nanoparticles between AD and control subjects. However, the lack of a relation between SIRM_100*K*_ and pathology does not contradict previous studies showing iron-oxides particles such as magnetite in the core of amyloid plaques^16^, but it shows that, if a biological mechanism is able to accumulate or precipitate magnetite nanoparticles in the brain during AD pathology, these changes are only minor and SQUID magnetometry would not be sensitive enough to detect them.

In summary, the results of this study showed good replicability and with our earlier work. Furthermore, we note that our current study was performed on a different SQUID magnetometer than earlier, thus supporting the inter-site repeatability of this methodology. We believe that the impact of repeatabiliy and replicability should not be underestimated in these type of experiments, especially given the SNR at high temperature.

Finally, we investigate to what extent SQUID magnetometry can detect formalin-induced changes in the magnetic carrier of the brain. Formalin fixation works via the reaction of formaldehyde with water to form methylene glycol, leading to low-concentrated free formaldehyde in the fixation solution^30^. Following a rapid penetration of the solution into the tissue, a slower process of protein cross-linking occurs. The reduction of the tissue water-content during fixation is only weakly affected, reaching 1% in the temporal cortex^30^. The question of whether formalin fixation introduces a bias in the magnetic analysis due to a chemical/structural transformation of the iron, has been sporadically investigated. The vast majority of the literature focuses on total iron levels, as assessed by atomic absorption spectroscopy and inductively-coupled mass spectrometry. One study performed on paired samples with and without formalin-fixation, over a much longer period of time than the current study, has shown that iron concentration decreased during fixation in 7 of the 8 investigated samples of neocortex; the amount of iron loss (43 %) did not change between white and grey matter; and metal leaching depended on the fixation period^45^. However, particles leaching out or dissolving from the tissue may behave very differently than protein-bound or labile trace elements, since magnetite is insoluble in water^46^ and requires an acid environment to be dissolved: a situation that was not encountered in this study.

One magnetometry study performed on brain tissue observed a reduction of ~ 45% in the SIRM as a result of one week of formalin fixation^22^, but in contrast, another study on human liver and spleen reported no evidence of chemical transformation of the iron after processing of the tissue, based on Mössbauer spectroscopy^23^. However, the latter publication reported that the ratio of the spectral areas of the two Mössbauer doublets was found to differ significantly between the fixed and frozen samples; a difference which they ascribed to inhomogeneity in the heme:non-heme iron ratio throughout the sample. In addition, the authors found a consistent leakage of the iron from the tissue up to 3% in the first 60 days, but after this, no further leaching was detected. Another study of Mössbauer spectroscopy on brain material reported the appearance of a doublet only in the low-temperature spectra of the formalin-fixed tissue, which was interpreted as iron leaching out of ferritin and hemosiderin, and bound to a non-specified chelating agent^47^.

Our study shows that SQUID magnetometry did not find a strong effect of formalin in dissolving either magnetite or ferrihydrite. To investigate the effects of formalin, a Bland-Altman plot was constructed. This method represents essentially a measure of repeatability from the within-subjects standard deviation of the replicates^48^, under the assumption that the formalin-fixed sample yields the same SIRM as the frozen equivalent. From these plots we can conclude that the limits of agreement are quite broad and include most of the data points, suggesting that SQUID magnetometry may not have enough sensitivity to detect changes in magnetic carriers induced by formalin fixation. Additionally, it appears that the scatter around the bias is larger as the average gets higher. This further suggests that in those samples containing a larger amount of magnetic carriers, a change due to fixation could be present and detectable, although the effect is so minor that it did not yield statistical significance upon testing. The reason behind the large standard devation of SIRM values may be, in addition to stocastic measurement error, that the paired samples are effectively different tissue blocks, which could contain a different, although small, fraction of white matter. This last issue could be quite important as the iron, ferritin, and water content of the grey matter are known to differ significantly from the white matter^49–51^. Therefore we can conclude that the change in SIRM values due to fixation, if true, would be less than or equal to the intra-subject variation, and definitely less than the precision of the technique.

Finally, we acknowledge that a longer fixation period may affect SIRM values to a larger extent than reported here, therefore minimal formalin storage (i.e. equal to or less than one year) is advised. We also foresee that SQUID magnetometry would be sensitive to detect possible changes induced by the transformation/leaching of iron particles due to formalin-fixation, if diseases characterized by extreme tissue iron accumulation were to be investigated. Therefore, the use of formalin-fixed material as a starting point for magnetometry studies should be evaluated on a case-by-case basis.

## Acknowledgments

This work was supported by the Netherlands Organization for Scientific Research (NWO) through a VENI fellowship to L. B (016.Veni.188.040). Stuart Gilder is thanked for useful discussion, Laurens Boers is thanked for help in sample preparation. Ekkes Brück is thanked for providing access to the SQUID magnetometer. The Netherlands Brain Brank and Laura Jonkman are thanked for providing the material for this study.

## Author contributions statement

L. B., A. W. and L.v.d.W. conceived the experiments. L.B. and A. L. conducted the experiments. L.B. and R. E. analysed the results. L.B wrote the paper and all authors read and contributed comments to the work.

## Additional information

The authors declare no competing interests.

